# Revisiting annotation of *Schistosoma mansoni* Micro-Exon Gene (MEG) family

**DOI:** 10.1101/2023.06.21.545858

**Authors:** Stepanka Nedvedova, Davide De Stefano, Olivier Walker, Maggy Hologne, Adriana Erica Miele

## Abstract

Genome sequencing of the human parasite *Schistosoma mansoni* revealed an interesting gene superfamily called *micro-exon gene* (MEG) that encodes MEG secreted proteins. The genes are composed of short exons (3-81 base pairs) with symmetrically inserted long introns (up to 5 kbp). This article recollects 35 *S. mansoni* specific *meg* genes that are distributed over 7 autosomes and one pair of sex chromosomes and that code for at least 87 verified MEG proteins. We used various bioinformatics tools to produce an optimal alignment, propose a phylogenetic analysis and highlight intriguing conserved patterns/motifs in the sequences of these MEG proteins. Based on the analyses, we were able to classify the MEG proteins into two subfamilies and to hypothesize their duplication and colonization of all the chromosomes. Together with motif identification, we also proposed to revisit MEGs’ common names and annotation in order to avoid duplication, to help reproducibility of research results and to avoid possible misunderstandings.

**Author Abstract:** *Schistosoma mansoni* is a parasitic worm, the etiological agent of schistosomiasis or bilharzia, a chronic tropical disease. It is a vector-borne parasite with a complex life cycle and an equally complex genome, assembled in 7 autosomes and a pair of sexual chromosomes. Within the gene products, one superfamily is particularly interesting, since it is specific to *Schistosomatidae*, highly variable and redundant: the micro-exon gene (MEG) family. As the name implies, these genes are made by short coding exons (3 to 81 base pairs), symmetrically interspersed by long introns (from 0.2 to 5 kbp). There are 35 *megs* allover the chromosomes, which code for at least 87 MEG proteins. We have aligned all of them, constructed a phylogenetic tree and proposed a theory for their duplication and genome colonization. Based on that, we propose a rational nomenclature to help the community to study MEG’s elusive role. We also propose to help WormBaseParaSite to adopt this new nomenclature to avoid giving the same acronym to different protein sequences.

## Introduction

Global efforts for genome sequencing after the human genome project [1,2] have been extremely beneficial to the entire scientific community. The Wellcome Trust Sanger Center, among others, undertook the task of sequencing the genomes of pathogens, in particular parasites, which affect the vast majority of the world population [3,4]. Parasites are complex eukaryotes, whose genomes show more resemblance with their hosts than with non-parasitic family members. In the last twenty years the completion of parasites’ genomes opened the way to study the molecular basis of the diseases caused by these agents and also to find and validate new drug targets. This last point is a critical one since most of the drugs against parasites are old, most of the time with severe side effects, or not anymore effective due to resistance. Nevertheless, focussed genetics and molecular biology studies had already been carried out before the genomic era, therefore quite a number of gene products had been annotated and served as landmarks for the recent annotations.

Under many respects, *Schistosoma mansoni*, the widest-spread agent of human schistosomiasis, is a good case study [4,5]. This vector-borne extracellular parasite is a digenean trematode, therefore metazoan, it possesses seven autosomes and a pair of sexual chromosomes, with a complex genetic structure including long interspersed sequences, transposon-like sequences, alternative splicing and gene duplications [5–9].

One class of genes in particular has attracted the attention of researchers, the so-called Micro-Exon Genes (MEG). Their discovery happened well before the completion of *S. mansoni* genome sequencing [10,11], although their annotation was completed with the latest version available on WormBaseParaSite (WBPS17 – WS282, accessed on March 2023) [12]. In total, 35 genes are annotated with characteristic symmetrical alternants of long introns (ranging from 0.1 to 5 kbp) and very short exons, whose length can vary from 6 to 81 base pairs, with a majority of 15 bp-long exons [5,12]. The introns’ structure is also quite intriguing and characteristic: their length is shorter towards the 5’ and 3’ ends (ranging from 100 to 500 bp), and longer (up to 4 kbp) in the center. This peculiarity made their automatic inference complicated from a genomic point of view.

However, the last fifteen years have seen plenty of transcriptomics and proteomics studies accumulating evidence of MEG complexity and variability and allowing their tracing back to the genome [5,11,13–16].

In this work, we present a systematic study aiming at understanding the putative origins and spreading over the entire genome of the MEG superfamily of genes. We also suggest a revision of the nomenclature in order to resolve some ambiguities in the literature.

## Methods

We have interrogated WormBaseParaSite (WBPS) database [12] over the last two years and we present the data from the latest session WS282 (accession 31^st^ March 2023), searching, within the genome of *Schistosoma mansoni,* for the terms “Micro-Exon Gene”, “MEG”, “antigen 10.3”, “GRAIL”, “ESP”. Per each gene, WBPS presents its structure, the possibility to retrieve the sequence, the position on the chromosome, and a translation identified with (at least) one associated UniProt ID. An example is given in Figure 1.

**Figure 1.**
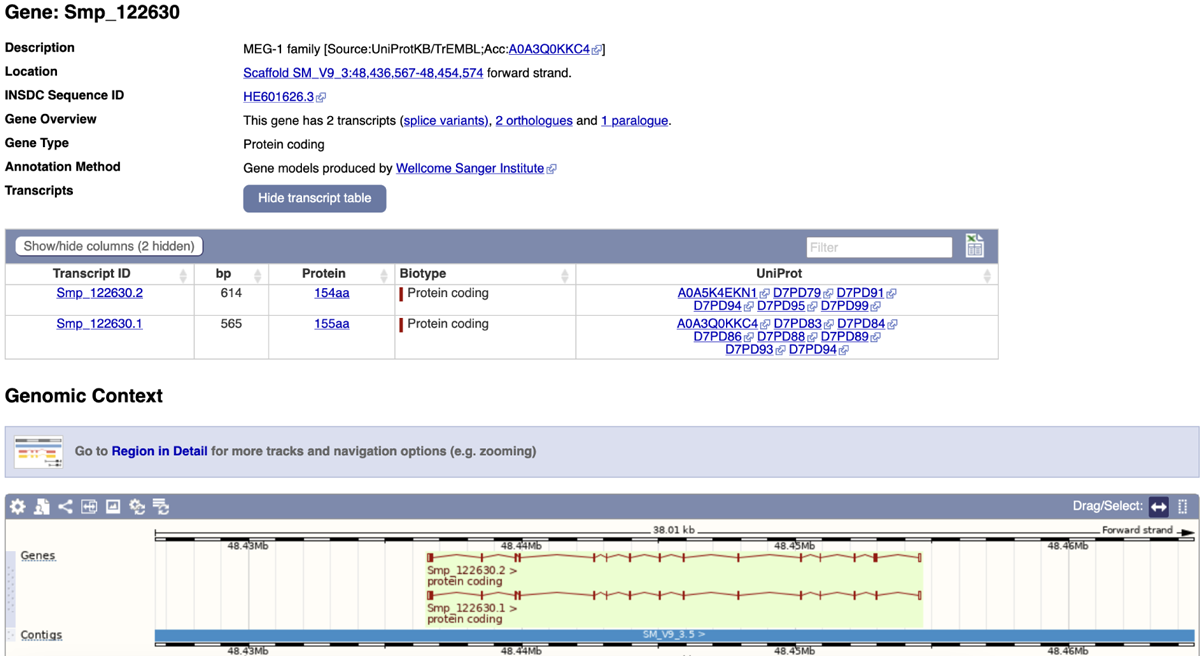
Screenshot of the entry for MEG-1 family on WormBaseParaSite [5].

In parallel, we have performed exactly the same search on UniProt KnowledgeBase [17] and cross-verified the results. All the protein sequences downloaded from UniProt were passed into psi-BLAST from NCBI [18], restricting the search to *S. mansoni*, in order to collect the maximum number of protein sequences annotated as MEG.

Afterwards, we performed a trimming, eliminating the duplicated entries, which displayed the same sequence with two different accession numbers, one from NCBI and one from UniProt. Whenever this was the case, we arbitrarily kept the UniProt ID only, for consistency with the cross-annotation of WBPS.

Primary structure alignment of MEGs was performed with T-Coffee [19], K-align [20] and MUSCLE [21] on the EBI server [22]. All the default parameters were chosen. Alignments were then manually inspected and finally the one by MUSCLE was preferred for subsequent analysis (Supplementary Figure S1) because the inserted gaps respected the exons’ boundaries; hence they better respected the biological constraints of alternative splicing.

Phylogenetic trees were built with both Simple phylogeny [23] and PRANK [24] on the EBI server [22]. The one from Prank was more consistent with the gene clustering and also with the type of retrotransposon sequences which had been found at the boundaries of the *megs* [7,8]; therefore it was retained and visualised on iTOL (interactive Tree of Life) server [25].

Emboss on the EBI server [22] was used to put in evidence conserved linear motifs, which were displayed with Weblogo [26].

Primary sequence analysis was performed by the ProtParam tool [27] on the Expasy website [28]; the results on calculated molecular weights, isoelectric point, aliphatic index and GRAVY index are presented in the supplementary Table S1.

## Results and Discussion

The latest annotated genome version of *S. mansoni* contains a total of 35 unique micro-exon genes (*meg*) with a peculiar symmetrical structure of 10 to 20 very short exons interspersed with long introns, whose length spans from 100 to almost 5000 base pairs. Unsurprisingly, the automatic annotation was challenging to trace these genes and to recognize them as protein expressing ones. The short exons are in the majority of cases a multiple of 3 and range from a minimum of 6 bp to a maximum of 81 bp. However, in a few cases one or two exons contain a number of base pairs not divisible by three. In the vast majority, the exons are 15 bp long, thus coding for 5 amino acids [5,11,12].

These 35 genes are interspersed over the 7 autosomes and the sexual chromosomes (Figure 2); most are coded on the leading strand and a few in the complementary one, such as *Smp_010550* (uncharacterized/MEG-15), which on the leading strand codes for one tRNA. On chromosomes 1, 3 and 5 they happen to be clustered together, reinforcing the hypothesis of their origin by gene duplication.

**Figure 2.**
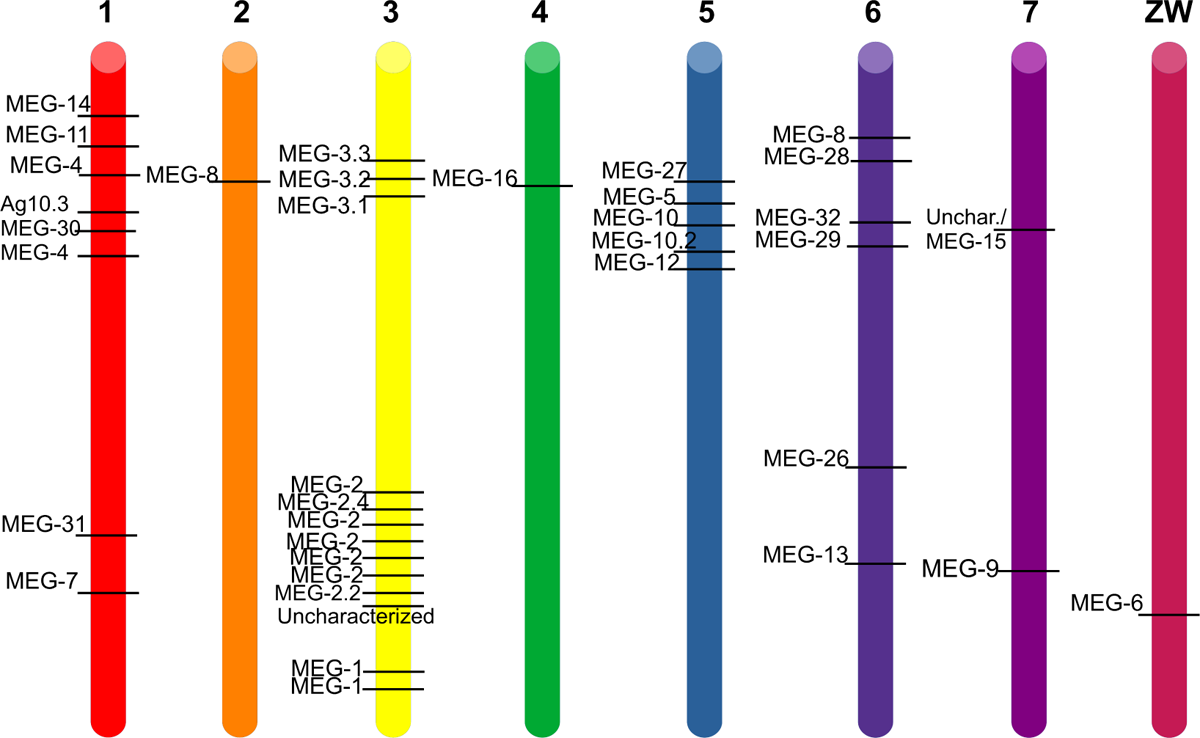
Schematic representation of S. mansoni haplotype. The approximate position of each MEG gene on each chromosome (coloured cylinder) is indicated by a black bar and its name on WormBaseParaSite is noted on the left. The chromosome number is on top of each cylinder.

All the chromosomes contain at least one *meg* (chromosomes 2, 4 and the sexual one), while chromosome 3 hosts thirteen distinct *meg*s. Moreover, close to the 5’ of *meg*s three types of (retro)transposable elements have been found, suggesting a spreading by gene duplication and transposition, and subsequent mutation. This finding was previously corroborated by studying the ratio between non-synonymous over synonymous (dN/dS) mutations in *meg* [7,8,29].

### MEG filiation

In *S. mansoni,* 35 *meg*s code for at least 87 verified MEG proteins with a unique UniProt ID, mainly originated by alternative splicing of the central exons. In the past, the gene sequences have been annotated and clustered into 23 families, numbered from 1 to 16 and from 26 to 32. MEG proteins have been found prior to the genome sequencing, in the secretions from eggs and adult worms, both in the transcriptomics and proteomics studies. At the beginning, they were named “antigen 10.3”, “Egg secreted protein no 15 (ESP15)”, “Grail”, before their peculiar gene structure was discovered [5,10,11,13,14,30].

We have aligned the 87 verified proteins, whose sequences are given in the supplementary Table S1, and we found that, among the three most used softwares offered by the EBI tools webpage [22], MUSCLE [21] was the one that reflected the most the splicing constraints, in fact it inserted the gaps between exons and not inside (supplementary figure S1), and it aligned the N-termini and the C-termini better than T-coffee or K-align.

Starting from an unknown ancestor, possibly on chromosome 6, a putative (phylogenetic) ontogenetic tree based on sequence similarities was built by PRANK [24] and a clustering in two main groups/clades was highlighted (Figure 3).

**Figure 3.**
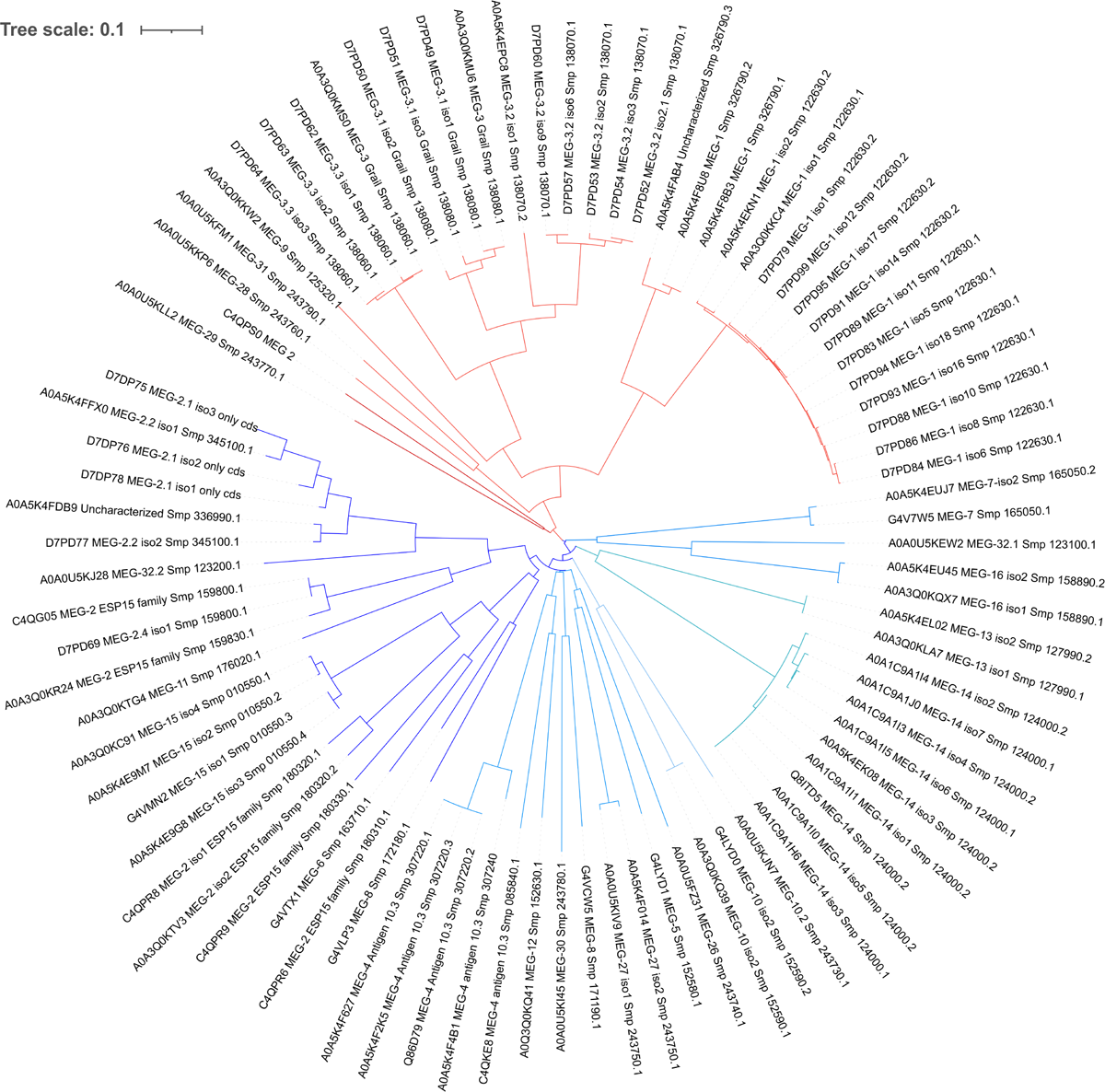
Phylogenetic tree coloured in red and blueafter clustering of the clades by sequence similarity. In the red clade an early event has separated MEG-29 and MEG-2 (ESP15, coded by Smp_183040.1) from the rest, so we have coloured this branch in dark red. Similarly, on the blue clade, MEG-7, MEG-32 and MEG-16 departed early from the clade and are highlighted in light blue.

It is plausible that a “proto-*meg*” was on chromosome 6, since the first leafs departing from each clade are there. Indeed MEG-28 and MEG-29 of the red clade and MEG-8 (*Smp_172180.1*) and MEG-32 of the blue clade are on chromosome 6. Of course this is just a pure hypothesis based on sequence similarity among the orthologous proteins.

The proposed filiation of the red clade, composed of 7 gene families, indicates an early jump on chromosome 3 to give rise to the MEG-1 and MEG-3 protein families, which practically colonised the entire chromosome. On the other hand, *meg-9* and *meg-31* jumped, respectively, on chromosomes 7 and 1. Based on sequence similarity, an event of gene duplication to generate *meg-3* and *meg-9* prior to splitting into two different chromosomes could be hypothesized, while *meg-31* could well be a filiation of *meg-28*.

Again on chromosome 6, we find the genes coding for MEG-13, MEG-8 (*Smp_172180.1*), MEG-26, MEG-32 of the second clade, the blue one in figure 3. Therefore, we might speculate that the first “experiments” on increasing protein variation by gene duplication and alternative splicing were carried out in chromosome 6 and then pursued on other chromosomes, in particular no. 3 and no. 1. In fact the protein coded by *meg-8* on chromosome 6 is close to the proteins of the MEG-2 family of chromosome 3, which then expanded and duplicated. The goal of increasing variability is quite common to parasites and it is usually linked to strategies for achieving host immune evasion [9,11,29,31,32].

*meg-9* on chromosome 7 was “joined” by *meg-15*, whose sequence is similar to the large *meg-2* protein family; hence we might speculate a jump from chromosome 3 to no. 7, rather than an intra-chromosomal duplication and mutation. The fact that they are not clustered together might corroborate this hypothesis.

Chromosome 3 is particularly interesting since it contains the highest number of *meg* genes and also the majority of egg-secreted MEG proteins (MEG-2, MEG-3, and MEG-1) [5,14,30] as well as the genes with the highest number of spliced isoforms (14 MEG-1 isoforms from Smp_122630, Figure 1).

According to genome annotation, *meg-8* (Smp_163710.1), *meg-16* and *meg-6* are the only representatives on chromosomes 2, 4 and the sexual one, respectively. They all belong to the blue clade, but the protein product of *meg-16* shares more similarities with MEG-32, which may suggest an early jump from chromosome 6 to 4, while MEG-6 proteins are closer to the MEG-2 protein family, suggesting a transposition from chromosome 3 towards the sexual chromosome. Moreover, MEG-8 protein shares 50% similarity with MEG-30 suggesting a transposition from chromosome 1 to chromosome 2. It is worth reminding that the mRNA of MEG-6 and MEG-8 are highly present in the eggs and oesophageal excreted/secreted transcripts, and the respective proteins have been found in proteomic studies [13,14,30].

Looking at the tree in Figure 3, we could also speculate that the clade painted in blue was more successful in diversifying the sequences, an aspect that we have tried to highlight with three shades of blue. On the other hand, we could speculatively infer that the clade painted in red was more successful in implementing the alternative splicing to achieve diversity.

### Conserved motifs

Going back to the total alignment in Supplementary Figure S1, it is clear that there are very few consensus motifs conserved, except the N-terminal signal peptide, which is needed for secretion. We have therefore decided to present this alignment more graphically by using WebLogo [26] in Figure 4. Apolar residues such as Pro, Phe, Leu are regularly spaced and highly represented. Charged residues, mostly basic Lys, are more conserved in the C-terminal part, while glutamic acid (E) is interspersed at more or less regular intervals of 15-20 aa (without taking into account the gaps). Soon after the signal peptide, all the MEGs present a stretch of hydrophobic and aromatic residues ended by a basic one: FL*χπχπ*F*X*_6_W*p*(K/H/R). Where *X* is any residue, *p* is a polar amino acid and *χπ* is a hydrophobic one.

**Figure 4.**
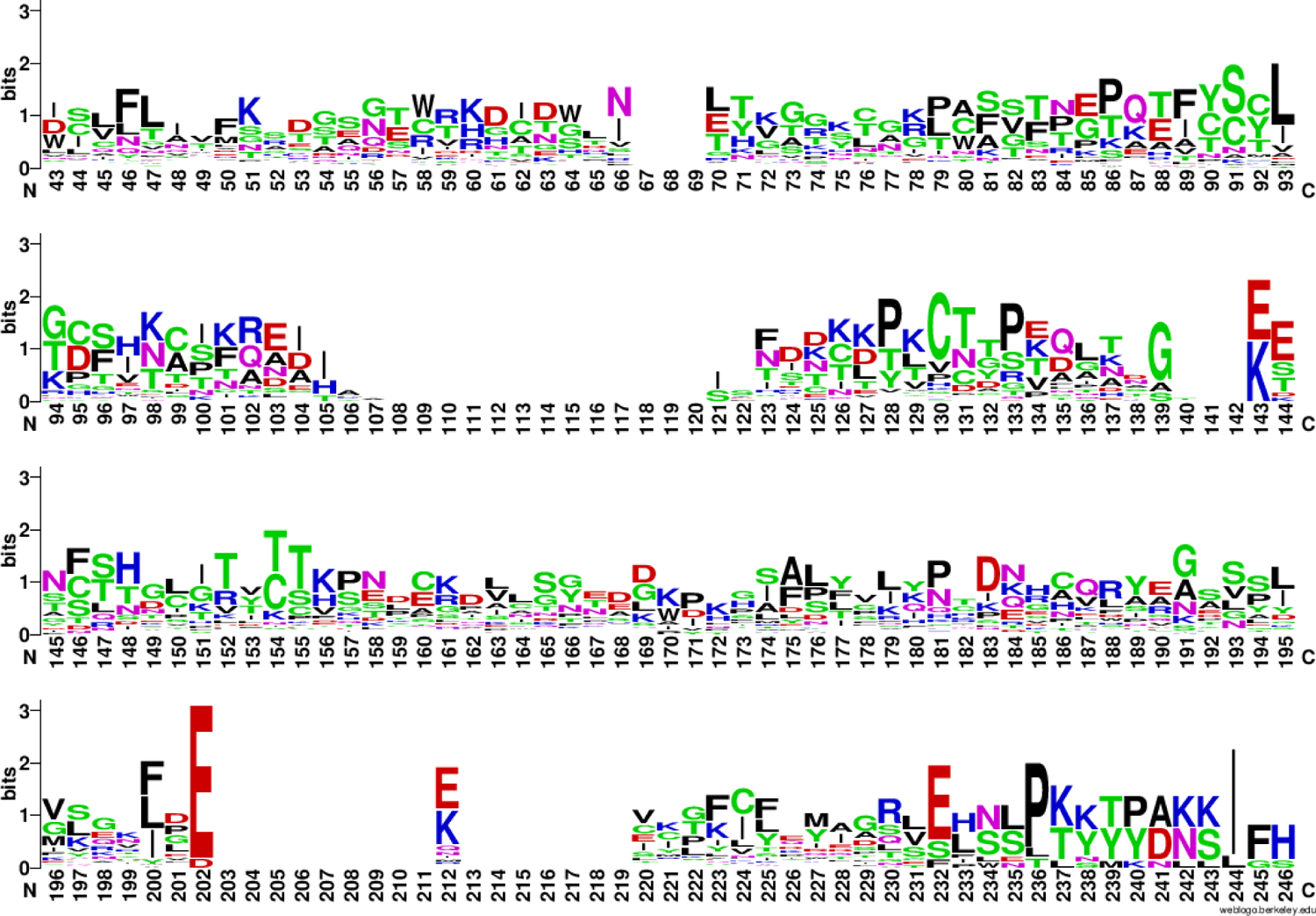
WebLogo representation of the alignment of all the 87 MEG protein sequences. The sequences of the signal peptide have been omitted for clarity. Even if the longest protein is 189 residues long, the number of gaps lengthens the aligned sequences to 246 residues.

Independently of the clade, it is apparent that MEG proteins possess sticky sequences, which we have verified by calculating the aliphatic index and the GRAVY index with ProtParam [28], data that we have included in the supplementary Table S1.

The stickiness of MEG proteins also appeared from the studies aimed at producing the isolated proteins by protein engineering. For example, recombinant MEG-3.2 and MEG-3.4 were purified from the inclusion bodies of *Escherichia coli* Rosetta cells and refolded by dialysis after purification, before they could be used for immunization studies [33]. MEG-14 was poorly expressed in bacteria and its amphipathic character was studied by circular dichroism with synchrotron light [34], before using the protein for binding studies with host factors [35]. MEG-4 (Sm10.3) was expressed in *Escherichia coli* Rosetta cells and purified in a buffer containing 0.5 M NaCl [36], a non-physiological hypertonic concentration of salt. MEG-24, MEG-27 and MEG-2.1 could not be produced in a heterologous host and, given their short size, they were chemically synthesized to perform *in vitro* studies [37, Nedvedova et al., submitted]. Chemical synthesis was also employed to use peptides of several MEG proteins as baits in search for host partners; this was the case for example of MEG-12 [32], MEG-8 [38] and twelve other MEG proteins expressed in the tegument and oesophageal glands [39].

If we split the two clades and align the sequences separately (35 proteins for the red clade and 52 for the blue clade), we can appreciate that the contribution of conserved Cys and Phe to the overall alignment comes from the red clade (Figure 5). On the other hand, the basic residues at the C-termini are contributed by the isoforms of the blue clade. Proline residues are conserved in both clades.

**Figure 5.**
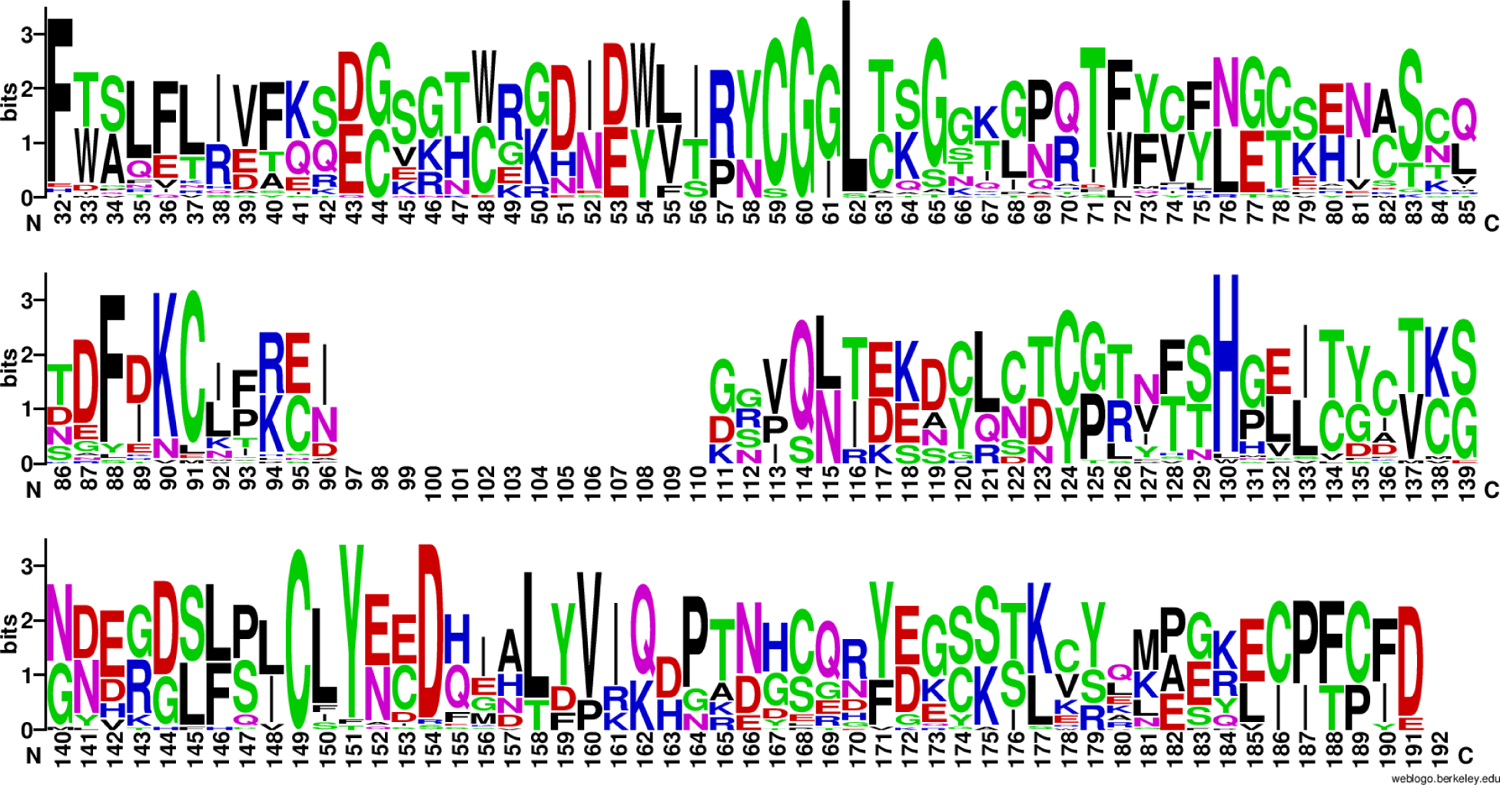
WebLogo representation of the alignment of the red clade composed of 35 MEG proteins, i.e. isoforms of MEG-1, MEG-3, MEG-9, MEG-28, MEG-29, MEG-31 and C4QPS0 of MEG-2 family. The sequence of the signal peptide has been omitted for clarity.

A hydrophobic motif at the N-terminus, soon after the signal peptide, is also present in the red clade (Figure 5), but with a slightly different sequence [FxxLFL(I/R)(V/D/E)Fxx(D/E)]. Moreover, we can appreciate that this first linear motif is followed by other four conserved motifs: CGGL *pp*G; (D/E)F(D/I/E)KCøø(R/K); C*x_5/7/9_*H*x_3/5/7_*C; and CLY*pp*D*X_3_*L(Y/F/D)V. In total, five short linear motifs characterize the red clade from the N- to the C-terminus, the first one being in common with the blue clade. It would be interesting to experimentally check whether these peptides are conserved because they are antigenic or because they confer some structural features to the IDPs.

### Nomenclature

Based on this classification and filiation, we would like to propose a more rational annotation of the gene products, trying to eliminate the gap between MEG-16 and MEG-26, and also possibly to rationalise the nomenclature of the large MEG-2 family, whose gene products have been numbered somehow arbitrarily. It is worth mentioning that this class possesses at least 9 more members deposited on UniProt, which we have excluded because they have been found only as mRNA, not yet as protein and there is apparently no gene associated to them in WBPS.

To start a reclassification we have taken the sequences of 13 MEG-2 proteins of the blue clade and aligned them (Figure 6), starting from the PRANK results. These proteins are coded by eight genes consecutively clustered together in the second half of chromosome 3 on the leading strand (Table 1). Only two non-meg coding genes (*Smp_326510* and *Smp_309120*), one after the second and one before the last *meg-2*, interrupt the chain.

**Figure 6.**
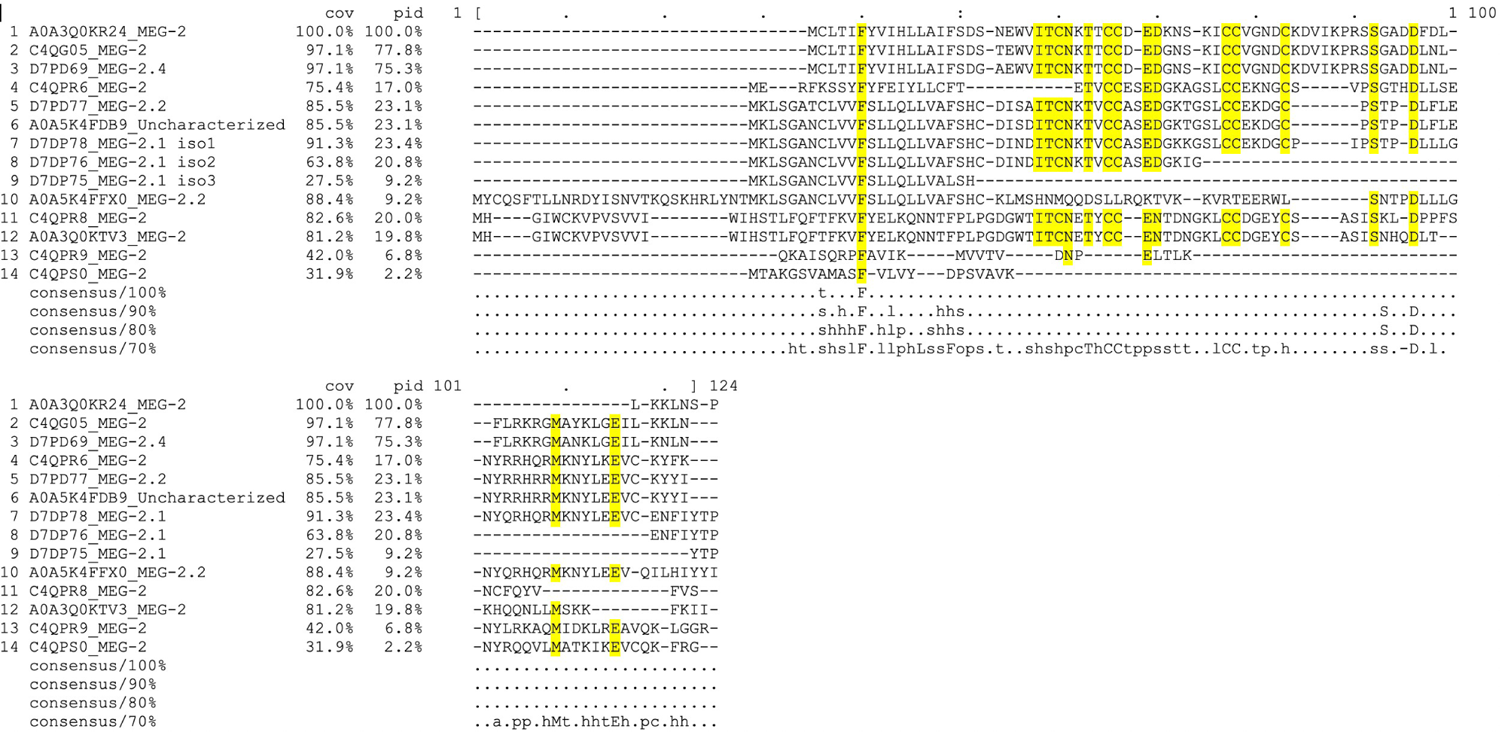
Sequence alignment of proteins coded by the meg-2 family. Sequences from 1 to 13 belong to the blue clade and the latter sequence (#14) belongs to the red clade. Conserved residues are highlighted in yellow.

**Table 1.**
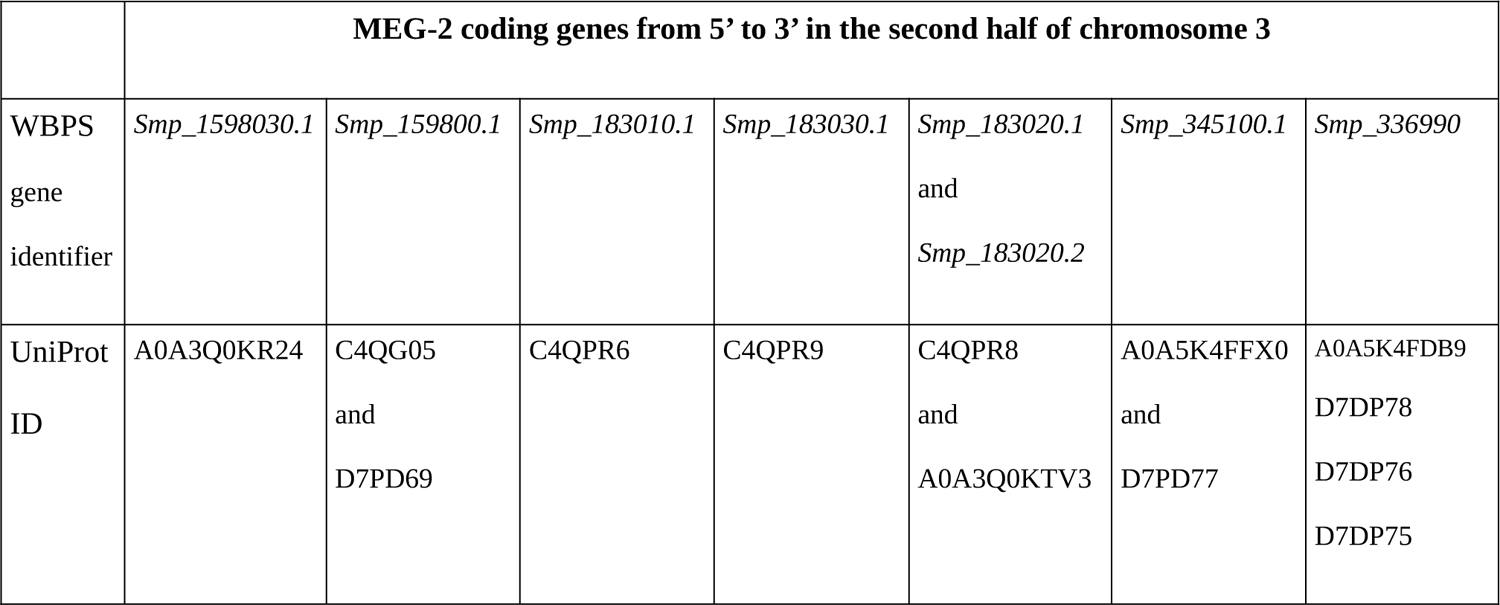
Summary of the cluster of the blue components of meg-2 on chromosome 3.

Interestingly, the gene coding for the red clade MEG-2/ESP15 isoform C4QPS0, *Smp_183040.1*, is located in the same part of chromosome 3, between *Smp_183010.1* and *Smp_183030.1*.

Based on ontology and genome positioning, we propose to keep the name MEG-2 to C4QPS0 of the red clade and to rename the products of the blue clade consecutively, according to their position on the genome, from 5’ to 3’, as indicated in Table 2 below.

**Table 2.** Proposed changes in the nomenclature of MEG protein products −2, −4, −8, −10 based on the filiation presented in *figure 3*.

| WBPS Gene identifier | UniProt identifier | Old name | New name |
| --- | --- | --- | --- |
| *Smp_183040.1* | C4QPS0 | MEG-2/ESP15 | MEG-2 |
| *Smp_1598030.1* | A0A3Q0KR24 | MEG-2/ESP15 | MEG-2.1 |
| *Smp_159800.1* | C4QG05 | MEG-2/ESP15 | MEG-2.2 isoform 1 |
|  | D7PD69 | MEG-2.4 isoform 1 | MEG-2.2 isoform 2 |
| *Smp_183010.1* | C4QPR6 | MEG-2/ESP15 | MEG-2.3 |
| *Smp_183030.1* | C4QPR9 | MEG-2/ESP15 | MEG-2.4 |
| *Smp_183020.1* | C4QPR8 | MEG-2 isoform 1 | MEG-2.5 isoform 1 |
| *Smp_183020.2* | A0A3Q0KTV3 | MEG-2 isoform 2 | MEG-2.5 isoform 2 |
| *Smp_345100.1* | A0A5K4FFX0 | MEG-2.2 isoform 1 | MEG-2.6 isoform 1 |
|  | D7PD77 | MEG-2.2 isoform 2 | MEG-2.6 isoform 2 |
| *Smp_336990* | A0A5K4FDB9 | Uncharacterized | MEG-2.7 isoform 1 |
|  | D7DP78 | MEG-2.1 isoform 1 | MEG-2.7 isoform 2 |
|  | D7DP76 | MEG-2.1 isoform 2 | MEG-2.7 isoform 3 |
|  | D7DP75 | MEG-2.1 isoform 3 | MEG-2.7 isoform 4 |
| *Smp_307220.1* | A0A5K4F627 | Developmentally regulated antigen 10.3 | MEG-4.1 isoform 1 |
| *Smp_307220.2* | A0A5K4F2K5 | Developmentally regulated antigen 10.3 | MEG-4.1 isoform 2 |
| *Smp_307220.3* | Q86D79 | Developmentally regulated antigen 10.3 | MEG-4.1 isoform 3 |
| *Smp_307240.1* | A0A5K4F4B1 | MEG-4/antigen 10.3 | MEG-4.2 |
| *Smp_085840* | C4KE8 | MEG-4/antigen 10.3 | MEG-17 |
| *Smp_172180.1* | G4VLP3 | MEG-8 | MEG-8 |
| *Smp_171190.1* | G4VCW5 | MEG-8 | MEG-18 |
| *Smp_152590.1* | A0A3Q0KQ39 | MEG-10 isoform 2 | MEG-10.1 isoform 1 |
| *Smp_152590.2* | G4LYD0 | MEG-10 isoform 2 | MEG-10.1 isoform 2 |
| *Smp_243730.1* | A0A0U5KJN7 | MEG-10.2 | MEG-10.2 |

Moreover, to disambiguate the MEG-4/antigen 10.3 proteins encoded by the genes *Smp_085840*, *Smp_307220* and *Smp_307240*, all on chromosome 1, we propose to keep the name MEG-4.1 to the protein products of the originally deposited gene (*Smp_307220*), to call MEG-4.2 the product of the closer relative *Smp_307240* and to call *Smp_085840*’s product MEG-17 because it is more similar to MEG-12 than to MEG-4.

Analogously, to disambiguate the two MEG-8 proteins, we propose to keep the name 8 to the gene *Smp_172180.1* present on chromosome 6, given the filiation described above, and to rename the other as MEG-18.

We also propose a slight modification of MEG-10 where 2 sequences hold the same protein name (Table 2), as well as for the MEG-3/Grail family, which is composed of three genes coding for a total of 15 isoforms (Table 3).

**Table 3.** Proposed changes in the nomenclature of MEG-3 protein products, based on the filiation presented in *figure 3*.

| **WBPS Gene identifier** | **UniProt identifier** | **Old name** | **New name** |
| --- | --- | --- | --- |
| *Smp_138060.1* | D7PD62 | MEG-3.3 isoform 1 |  |
|  | D7PD63 | MEG-3.3 isoform 2 |  |
|  | D7PD64 | MEG-3.3 isoform 3 |  |
|  | A0A3Q0KMS0 | MEG-3 Grail family | MEG-3.3 isoform 4 |
| *Smp_138070.1* | D7PD52 | MEG-3.2 isoform 2.1 | MEG-3.2 isoform 1 |
|  | D7PD53 | MEG-3.2 isoform 2 |  |
|  | D7PD54 | MEG-3.2 isoform 3 |  |
|  | D7PD57 | MEG-3.2 isoform 6 | MEG-3.2 isoform 4 |
|  | D7PD60 | MEG-3.2 isoform 9 | MEG-3.2 isoform 5 |
| *Smp_138070.2* | A0A5K4EPC8 | MEG-3.2 isoform 1 | MEG-3.2 isoform 6 |
| *Smp_138080.1* | D7PD49 | MEG-3.1 isoform 1 |  |
|  | D7PD50 | MEG-3.1 isoform 2 |  |
|  | D7PD51 | MEG-3.1 isoform 3 |  |
|  | A0A3Q0KMU6 | MEG-3 Grail family | MEG-3.1 isoform 4 |

An interesting case is the one of MEG-15, which in the latest annotation lost its name and became “uncharacterized”, therefore we propose to go back to its name, given its similarity with the MEG-2 family and with MEG-6 (see figure 2). Indeed the gene *Smp_010550* has 4 splice variants each one coding for one isoform, identified on UniProt. A summary is given in Table 4.

**Table 4.** Proposed changes in the nomenclature of MEG-15 protein products

| WBPS Gene identifier | UniProt identifier | Old name | New name |
| --- | --- | --- | --- |
| *Smp_010550.1* | A0A3Q0KC91 | Uncharacterized protein | MEG-15 isoform 1 |
| *Smp_010550.2* | A0A5K4E9M7 | Uncharacterized protein | MEG-15 isoform 2 |
| *Smp_010550.3* | G4VMN2 | Uncharacterized protein | MEG-15 isoform 3 |
| *Smp_010550.4* | A0A5K4E9G8 | Uncharacterized protein | MEG-15 isoform 4 |

Finally, we think that a bit of order in the MEG-1 family would also improve the readability (Table 5), although this is, together with MEG-3, one of the best annotated family. It is better to underline that there are other isoforms on UniProt that have only been inferred by transcriptomics and do not belong to any of the three deposited coding genes; therefore they are not included in this list, nor they have been used for the alignment.

**Table 5.** Proposed changes in the nomenclature of MEG-1 protein products

| WBPS Gene identifier | UniProt identifier | Old name | New name |
| --- | --- | --- | --- |
| *Smp_326790.1* | A0A5K4F8B3 | MEG-1 | MEG-1.1 isoform 1 |
| *Smp_326790.2* | A0A5K4F8U8 | MEG-1 | MEG-1.1 isoform 2 |
| *Smp_326790.3* | A0A5K4FAB4 | Uncharacterized protein | MEG-1.1 isoform 3 |
| *Smp_122630.1* | A0A3Q0KKC4 | MEG-1 isoform 1 | MEG-1.2 isoform 1 |
|  | D7PD83 | MEG-1 isoform 5 | MEG-1.2 isoform 2 |
|  | D7PD84 | MEG-1 isoform 6 | MEG-1.2 isoform 3 |
|  | D7PD86 | MEG-1 isoform 8 | MEG-1.2 isoform 4 |
|  | D7PD88 | MEG-1 isoform 10 | MEG-1.2 isoform 5 |
|  | D7PD89 | MEG-1 isoform 11 | MEG-1.2 isoform 6 |
|  | D7PD93 | MEG-1 isoform 16 | MEG-1.2 isoform 7 |
| *Smp_122630.2* | A0A5K4EKN1 | MEG-1 isoform 2 | MEG-1.2 isoform 8 |
|  | D7PD79 | MEG-1 isoform 1 | MEG-1.2 isoform 9 |
|  | D7PD91 | MEG-1 isoform 14 | MEG-1.2 isoform 10 |
|  | D7PD94 | MEG-1 isoform 18 | MEG-1.2 isoform 11 |
|  | D7PD95 | MEG-1 isoform 17 | MEG-1.2 isoform 12 |
|  | D7PD99 | MEG-1 isoform 12 | MEG-1.2 isoform 13 |

### Towards a Function?

The exact role of the MEG proteins is still unknown; their high copy number and their high variability make inferring tough. In recent years many “omics” studies have boosted the research on schistosomes, with the aim to find new drug targets, to develop more early and precise diagnostics and to implement a vaccine. A handful of studies on individual MEG proteins have been carried out, revealing their nature as intrinsically disordered proteins, without or with a morphing/chameleon behaviour. One morphing IDP is MEG-14, which is able to fold upon binding to negatively charged membranes or to calgranulin, a human S100 family member involved in inflammation [34,35]. MEG-24 (whose sequence was not deposited in any public repository) and MEG-27 were shown to bind to liposomes and to agglutinate red blood cells, possibly through the formation of amphipathic helices [37].

We have recently characterised by NMR three splice variants of *Smp_336990* of the MEG-2 family and confirmed their IDP nature, together with some interesting hairpin loops, which might confer some rigidity and act as a platform for interactions with host partners [Nedvedova et al., submitted]. Indeed MEG-2 family members possess the highest content of Cys residues (Figure 5) and a conserved N-terminal motif Cys_2_X_(6/8/10)_Cys_2_, which reminds either of a Zn-finger or a [2Fe2S] cluster. It remains to prove that MEG-2 proteins might be metal sensors or chelators. A clue might be the fact that they are present in the oesophagus’ secretions, where hemoglobin digestion occurs; this digestion releases high quantities of iron, potentially toxic. It is known that hemozoin is regurgitated [40], but maybe MEG-2 proteins could act as an iron cleaner, limiting the oxidative stress.

MEG proteins high variability and their high expression in the mammalian host have made the researchers think about a role in host immune evasion or modulation. This was the basis for using short synthetic peptides issued from more or less each MEG family as baits to fish IgG from infected mouse models. The most antigenic ones were then used as protective vaccine, although with low efficacy [39]. The question arises whether the full-length isoforms would be more protective, or whether the conserved motifs that we have highlighted could be a better strategy.

## Conclusions

The 87 protein products of *Schistosoma mansoni*’s 35 micro-exon genes (MEG) are elusive and interesting macromolecules, a Pandora’s box of tools to understand the complex behaviour of these fascinating and dangerous parasitic worms. We have presented a rationalization of the gene families, based on the sequence similarities, and proposed a renaming in order to avoid confusion and to help trimming what is known (the tip of the iceberg) from what still remains to study.

## Acknowledgments.

We would like to dedicate this paper to the memory of the late Dr Ricardo DeMarco who pioneered the field of micro-exon genes and dedicated his research to unravel schistosome’s biology.

## Conflict of interest

The authors declare that they are free from any competing interests.

## Funding acquisition

This research received funds from Improvement in Quality of the Internal Grant Scheme at CZU, reg. no. CZ.02.2.69/0.0/0.0/19_073/0016944, financed from the funds of Operational Programme Research, Development and Education, in the framework of ESF Call no. 02_19_073 for Improving the Quality of Internal Grant Schemes at Higher Educational Institutions in priority axis 2 OP.

## Authors contribution

AEM and MH conceived the research. SN acquired the funding. SN and DDS performed analysis. SN, DDS, OW, MH and AEM analysed the data. All authors wrote, read and edited the manuscript.

## Notes

### Competing Interest Statement

The authors have declared no competing interest.

